# Evaluation of the bacterial and fungal microbiome on the skin of *Anaxyrus boreas*, *Rhinella marina*, and *Xenopus laevis* and implications related to *Batrachochytrium dendrobatidis* susceptibility

**DOI:** 10.1101/2020.07.19.211102

**Authors:** Moamen M. Elmassry, Scot Dowd, Cynthia Carey, Michael J. San Francisco

**Affiliations:** Department of Biological Sciences, Texas Tech University, Lubbock, TX; MR DNA, Molecular Research LP, Shallowater, TX; Department of Integrative Physiology, University of Colorado, Boulder, CO; Honors College, Texas Tech University, Lubbock, TX

## Abstract

The skin and its microbiome are the first lines of defense against environmental stressors and pathogens. The symbiotic relationships between the host and the microbiome and within the microbiome are critical to host health. In frogs, this research area is lacking especially in the context of their skin infection with *Batrachochytrium dendrobatidis*. *B. dendrobatidis* is a major fungal skin pathogen to amphibians that has caused the extinction of hundreds of amphibian species populations. While some frog species are known susceptible to *B. dendrobatidis* infections, and others are resistant, we hypothesized that the skin microbiome of frogs plays a role in the prevention of *B. dendrobatidis* infections that is yet unknown. Therefore, in this work we have examined the bacterial and fungal skin microbiome of one species that is sensitive to *B. dendrobatidis* infections, *Anaxyrus boreas* (formerly: *Bufo boreas*), and the resistant species, *Xenopus laevis* and *Rhinella marina* (formerly: *Bufo marinus*). This was accomplished using tag-encoded FLX amplicon pyrosequencing (bTEFAP) of the 16S rRNA and ITS DNA regions. Our results showed that the bacterial and fungal skin microbiome of *A. boreas* and *R. marina* were more similar than to that of *X. laevis.* We found distinct patterns between the skin microbiome of *B. dendrobatidis* sensitive and resistant frog species. For the bacterial microbiome, the most abundant bacterial genus observed in all frog samples was *Microbacterium*. The resistant species had higher abundance in the genera *Pseudomonas*, *Sphingobium*, *Pedobacter*, *Variovorax*, *Morganella*, *Sphingomonas*, *Giesbergeria*, and *Agromyces*. In contrast, they had lower abundance of *Elizabethkingia* (formerly: *Flavobacterium*)*, Enterobacter*, *Ochrobactrum*, *Arthobacter*, *Stenotrophomonas*, *Shinella*, *Klebsiella*, *Aeromonas*, *Comomonas*, *Chitinophaga*, and *Rhodococcus*. Regarding the fungal microbiome, the resistant species showed higher abundance in the genera *Aspergillus*, *Cladosporium, Elaphomyces*, *Monascus*, *Tritirachium*, *Ceratostomella*, and *Claviceps*, while *Asterotremella*, *Trichosporon*, and *Malasezzia* genera were less represented in resistant species. We have also observed that in the resistant species, the skin microbiome had higher diversity in the fungal microbiota, but not in the bacterial microbiota. We speculate that those observed differences have implications in *B. dendrobatidis* susceptibility that are yet to be determined, possibly through competition for nutrients or the production of specific anti-fungal molecules.

**GRAPHICAL ABSTRACT:** 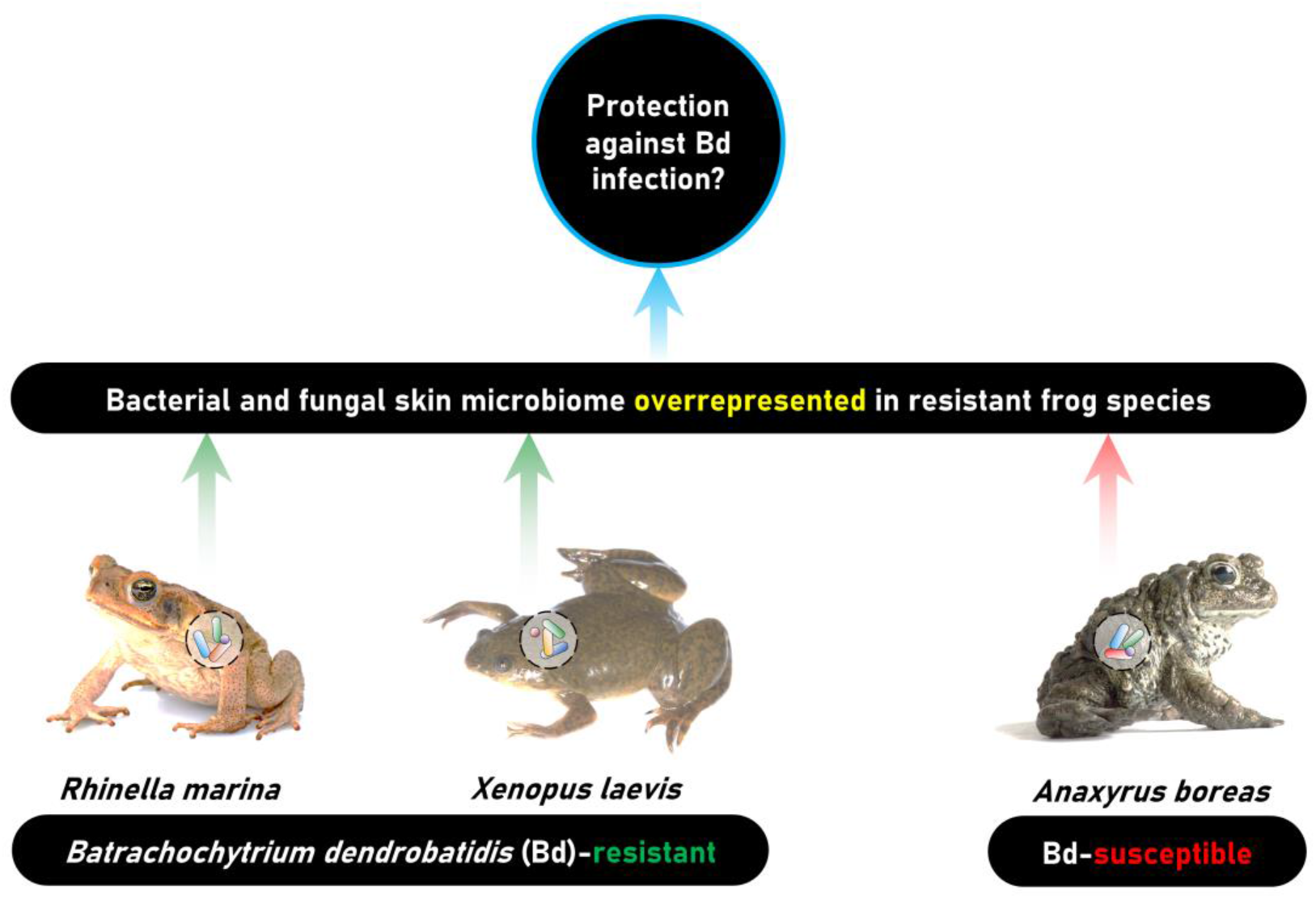

## INTRODUCTION

Global populations of amphibians have reached alarmingly low numbers due to the rapid spread of the chytrid fungus *Batrachochytrium dendrobatidis* (Bd) (Berger et al., 1998; Longcore, Pessier, & Nichols, 1999). The chytrid has the widest range of susceptible host species for any vertebrate pathogen known to date, and massive Bd-related die-offs have been reported in every continent inhabited by amphibians (Fisher, Garner, & Walker, 2009; Heyer & Murphy, 2005). The fungus causes skin infections that may result in chytridiomycosis and eventual death of the animals by disrupting ion homeostasis, a critical function of the skin of amphibians (Voyles et al., 2009). The recent distribution of infected hosts to geographic regions which lacked previous exposure to the pathogen (Fisher & Garner, 2007; Morse, 1995; Xie, Olson, & Blaustein, 2016). Most deaths caused by Bd infection occur in amphibians that breed in permanent bodies of water, suggesting that the pathogen is predominantly aqueous in nature (Lips et al., 2006). Infections are much more prevalent in the keratinized tissue of post-metamorphic stages than in pre-metamorphic stages, which may be the result of a pathogenic down-regulation of host immune response (Berger et al., 1998; Ribas et al., 2009; Rosenblum et al., 2009). A number of factors are thought to affect Bd susceptibility among amphibians, such as seasonal and global temperature change, and elevation (Berger et al., 2004; Bosch, Carrascal, Durán, Walker, & Fisher, 2007; Bradley et al., 2019; Kriger & Hero, 2006; Maniero & Carey, 1997; Muths, Pilliod, & Livo, 2008; Pounds et al., 2006; Raffel, Rohr, Kiesecker, & Hudson, 2006; Russell et al., 2019). Furthermore, other factors include previous exposure and sensitization to the pathogen, or a lack thereof, and individual skin microbiota (Cogen, Nizet, & Gallo, 2008; Culp, Falkinham, & Belden, 2007; R. N. Harris, James, Lauer, Simon, & Patel, 2006; Lam, Walke, Vredenburg, & Harris, 2010; Lauer et al., 2007; Lauer, Simon, Banning, Lam, & Harris, 2008; Morse, 1995).

A few studies have attempted to characterize the natural cutaneous microbiota of amphibians by culture-independent means (Abarca et al., 2018; Edwards, Byrne, Harlow, & Silla, 2017; Federici et al., 2015; Lauer et al., 2008; Madison, Ouellette, Schmidt, & Kerby, 2019). Cutaneous microbiota of certain amphibians are capable of producing antifungal compounds which may prevent Bd infection, and only a very limited proportion of resident host microbes can be identified with culture dependent methods (Abarca et al., 2018; Dowd, Sun, Secor, et al., 2008; Edwards et al., 2017; Federici et al., 2015; Lauer et al., 2008; Madison et al., 2019). In addition, a comparison of the cutaneous microbiota between resistant and non-resistant amphibians to Bd would be especially helpful in characterizing potential key players in Bd resistance. The aim of this study is to identify the natural skin microbiota of three species of frogs raised in a breeding facility: *Anaxyrus boreas* (formerly: *Bufo boreas), Rhinella marina* (formerly: *Bufo marinus*), and *Xenopus laevis.* While *A. boreas* is susceptible to Bd infections, *X. laevis* and *R. marina* are known to be resistant to Bd (Olson et al., 2013). Understanding the differences in the skin microbiome of these species is important as the skin microbiome is implicated in Bd susceptibility.

## RESULTS

### Bacterial microbiome of *A. boreas, R. marina*, and *X. laevis* skin samples

Twenty tagged and unlabeled samples of skin swabs from *A. boreas, R. marina* and *X. laevis* were distinguishable based on their representative bacterial microbiome. Samples from the same frog species clustered together (Fig. 1). Based upon the clustering algorithm analysis of the bacterial genera *A. boreas* and *R. marina* clustered together at a relative Manhattan distance of 1.5. The clustering distance based on bacterial microbiome between *A. boreas* and *R. marina*, and *X. laevis* was observed to be 3 (Fig. 1). But this could be due to the limited number of samples from *X. laevis.*

**Figure 1.**
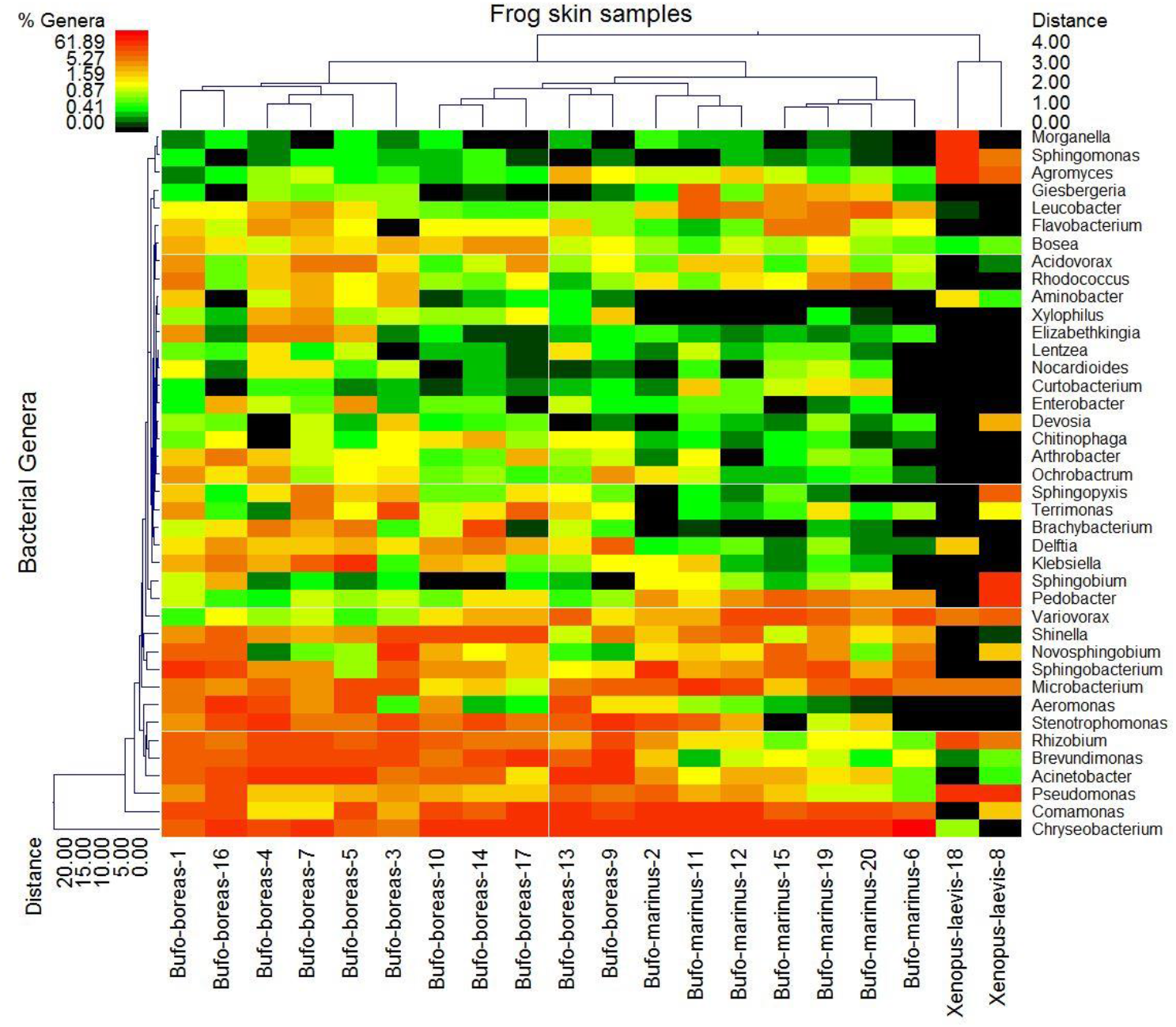
Double hierarchal dendrogram showing different bacterial genera abundance on the skin of *A. boreas*, *R. marina*, and *X. laevis*. Weighted-pair group clustering method, and Manhattan distance method with no scaling. The dendrogram linkages of the bacterial classes are not phylogenetic but based upon relative abundance of the bacterial classes within the samples. Clustering of the systems were similarly based upon abundance of the top 40 most abundant bacterial genera among individual samples. The heat map depicts the relative percentage of each genera of bacteria (Variables clustering on Y-axis) within each sample (X-axis clustering). The heat map colors represent the relative percentage of the bacterial genera within each sample with the legend indicated at the upper left of the figure. The samples along the X-axis with Manhattan distances are indicated by branch length and an associated scale located at the upper right of the figure. Clustering based upon Manhattan distance of the bacterial genera along the Y-axis and their associated scale is indicated in the lower left.

The most abundant bacterial genus observed for all three frog samples was *Microbacterium*. *Microbacterium* species are abundant in aquatic habitat, which explains its abundance in skin microbiome (Ashcroft, Zalinger, Bevier, & Fekete, 2007; Takeuchi & Hatano, 1998). The most prominent distinctions between *X. laevis* and the other two species (*A. boreas* and *R. marina*) belonged to the genera *Pseudomonas*, *Sphingobium*, *Pedobacter*, *Morganella*, *Sphingomonas*, and *Agromyces.* Each of these genera of bacteria were more abundant in samples from *X. laevis* than either of the *A. boreas* or *R. marina.* The following genera were relatively higher in *R. marina* than in *A. boreas: Giesbergeria, Leucobacter, Pedobacter*, and *Variovorax.*

*Elizabethkingia* (formerly: *Flavobacterium* or *Chryseobacterium*), *Enterobacter*, *Ochrobactrum*, *Arthobacter*, *Sphingobacterium*, *Stenotrophomonas*, *Shinella*, *Klebsiella*, *Aeromonas*, *Comomonas*, *Chitinophaga*, and *Rhodococcus* species were in relatively low abundance in *X. laevis* samples compared to both *A. boreas* and *R. marina* samples. The greatest discrimination between the two species *A. boreas* and *R. marina* was observed for *Aminobacter*, *Aeromonas*, *Xylophilus*, and *Brachybacterium*, which were more abundant in *A. boreas* samples. *Pedobacter* was more abundant in the *R. marina* samples compared to *A. boreas*. Lastly, *Variovorax* was more abundant in *X. laevis* and *R. marina* skin samples than in that of *A. boreas*. Overall, *X. laevis* samples had lower diversity than *A. boreas* and *R. marina* samples.

### Fungal microbiome of *A. boreas, R. marina*, and *X. laevis* skin samples

The clustering algorithm analysis of the fungal genera shows that the three species cluster together at a relative Manhattan distance of 3. Six genera that were observed in *X. laevis* skin swabs were absent from all samples of the two species (*A. boreas* and *R. marina*) sampled. These included *Cladosporium, Elaphomyces*, *Monascus*, *Tritirachium*, *Ceratostomella*, *Claviceps*, and *Aspergillus* in decreasing order of abundance (Fig. 2). Notably absent from *X. laevis* and present in either *A. boreas* or *R. marina* were *Saccharomyces*, *Asterotremella*, *Trichosporon*, and *Malasezzia* species. The genus *Asterotremella* was absent in all *R. marina* samples, but the genus *Saccharomyces* was absent from all but one sample of *A. boreas*. Overall, *X. laevis* samples had higher diversity than *A. boreas* and *R. marina* samples.

**Figure 2.**
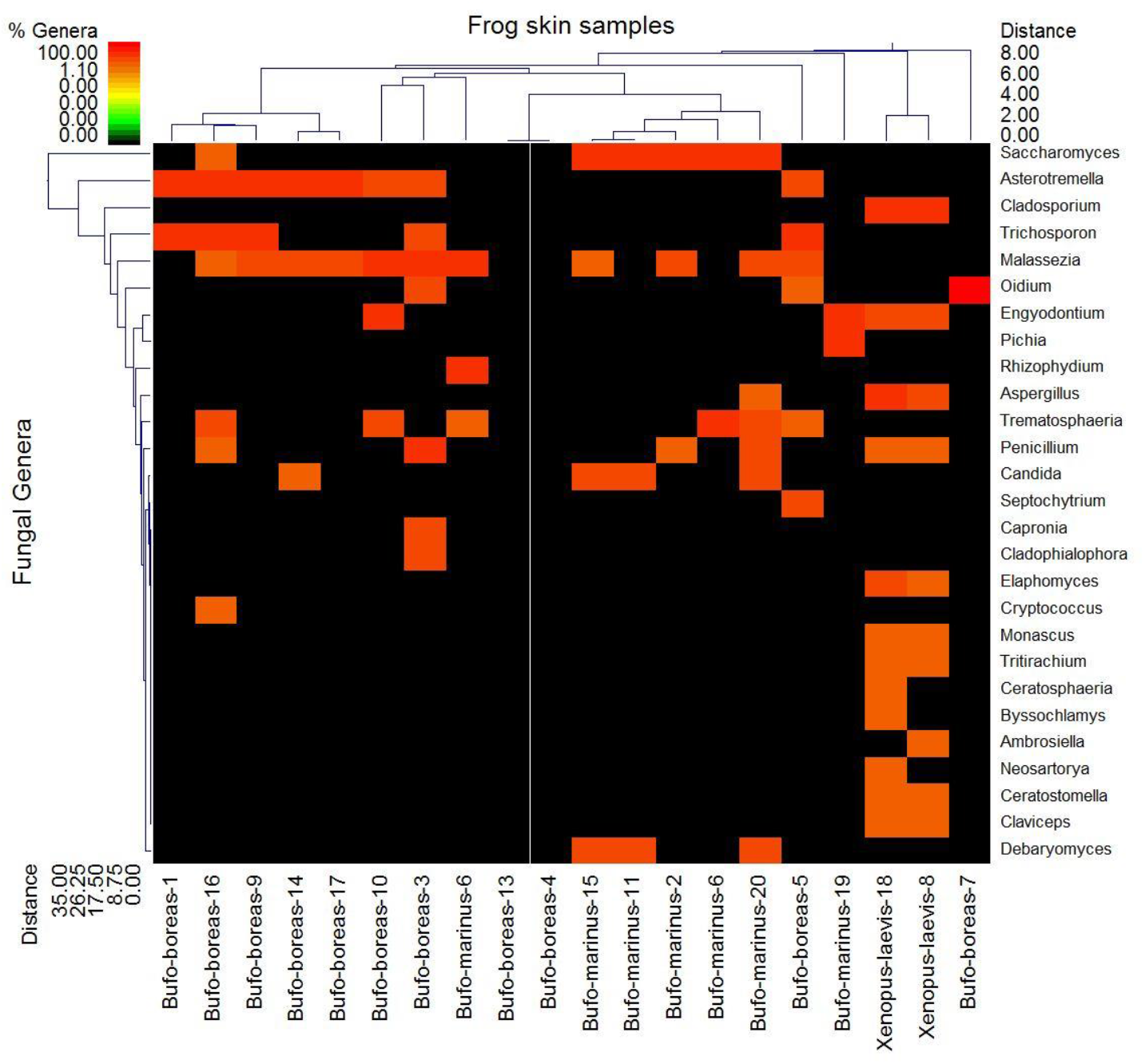
Double hierarchal dendrogram showing different fungal genera proportion on the skin of *A. boreas*, *R. marina*, and *X. laevis*. Weighted-pair group clustering method, and Manhattan distance method with no scaling. The dendrogram shows linkages of the fungal genera are not phylogenetic but based upon relative abundance of the fungal classes within the samples. Clustering of the systems were similarly based upon abundance of fungal genera among individual samples. The heat map depicts the relative percentage of each genus of fungi (Variables clustering on Y-axis) within each sample (X-axis clustering). The samples along the X-axis with Manhattan distances are indicated by branch length and an associated scale located at the upper right of the figure. Clustering based upon Manhattan distance of the fungal genera along the Y-axis and their associated scale is indicated in the lower left.

## DISCUSSION

The diversity of resident skin microbes from three naïve species of frogs was examined using tag-encoded FLX amplicon pyrosequencing (bTEFAP) with much greater detail than what was previously possible using traditional culture methods (Dowd, Wolcott, Sun, et al., 2008). Cutaneous microbiota is shown to be relatively unique to each of the three species, which clustered at a clear exclusion of one another for the bacterial and fungal microbiome analysis. Most interestingly, the Bd-resistant *X. laevis* is clearly distinct from the two Bd-susceptible species *A. boreas*, which have a notable degree of overlap in the genera of bacteria present on the skin. This suggests that the resident skin bacteria of *X. laevis* may contribute significantly to Bd immunity. However, further analyses with bigger population size are still needed to confirm our results.

Many of the bacteria detected on *X. laevis* are known to produce antimicrobial compounds, and some of these have even been shown to inhibit Bd growth, while the effect of the rest is still unknown (Table 1) (Culp et al., 2007; R. N. Harris et al., 2006; Lam et al., 2010; Lauer et al., 2007, 2008). *Pseudomonas* and *Pedobacter* genera that exhibit anti-Bd activity, were more abundant in the *X. laevis* samples than in those from *A. boreas* and *R. marina* (R. N. Harris et al., 2006; Lauer et al., 2007; Myers et al., 2012). It was also found that relative abundance of *Pseudomonas* was decreased with increasing Bd load in frogs that may hint to their competitive inhibition (Wilber, Jani, Mihaljevic, & Briggs, 2019). Another interesting observation is that both *A. boreas* and *R. marina* species have a ubiquitous and high abundance occurrence of *Chryseobacterium*, which has also been reported to possess anti-fungal activity (Lauer et al., 2007). Susceptibility to the chytrid fungus might persist in *A. boreas*, due to species-level variations within the *Chryseobacterium* genus. Furthermore, the genera *Pseudomonas* and *Chryseobacterium* were among the most prevalent taxa associated with Bd resistance (Jiménez, Alvarado, Estrella, & Sommer, 2019; Muletz-Wolz et al., 2017). The skin microbiome of *X. laevis* samples showed numerous bacterial genera such as *Sphingobium, Morganella, Agromyces*, and *Sphingomonas*, that haven’t been documented before whether they have the capacity to act against Bd, which is yet to be investigated. Lastly, these samples were obtained from “naïve” laboratory-raised frogs and their relative proportions may be different in the wild.

**Table 1.**
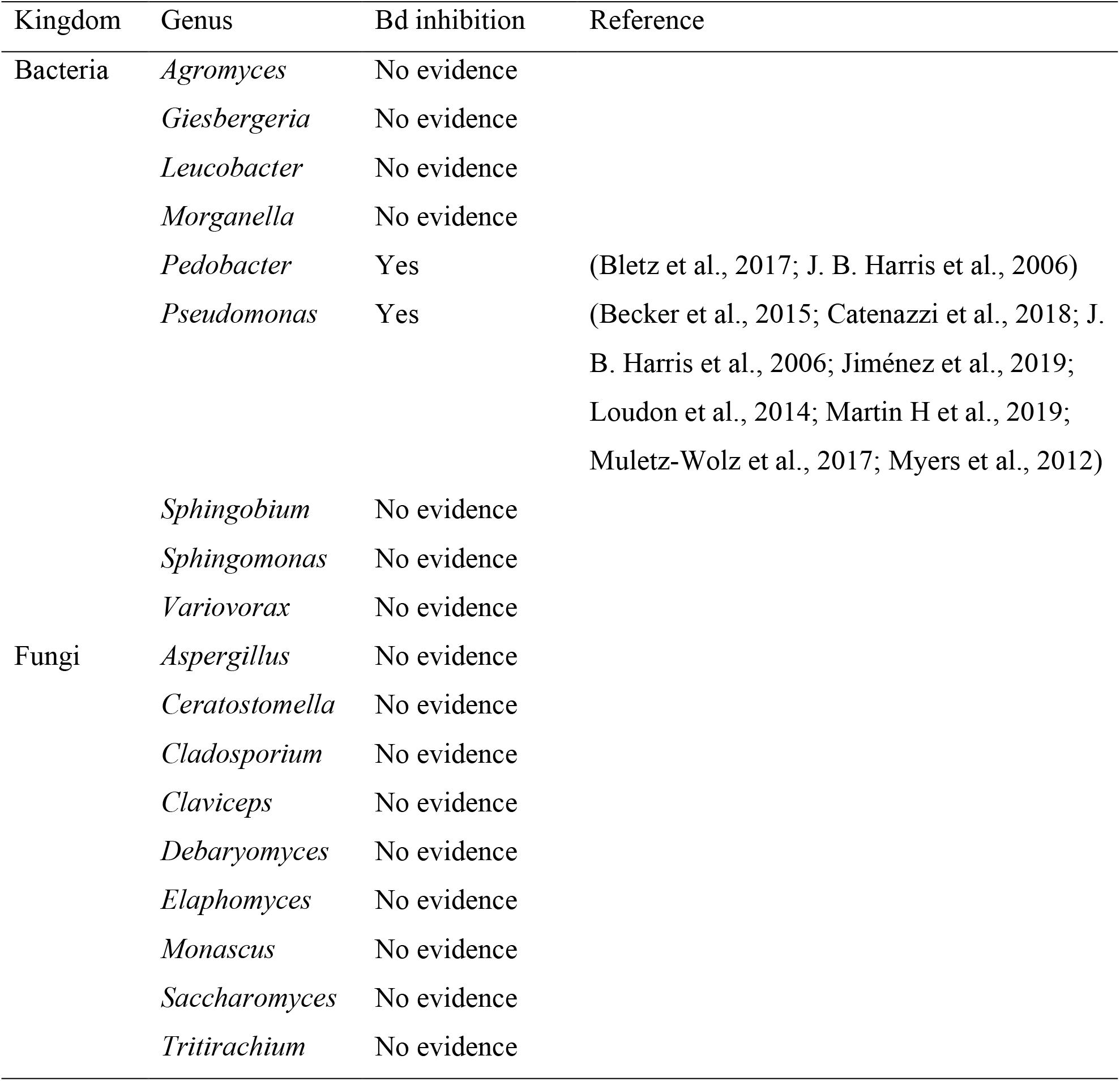
Bacterial and fungal genera that were overrepresented in the skin microbiome of resistant rather than sensitive frog species, and evidence supporting their *Batrachochytrium dendrobatidis* inhibition.

This is the first report of fungal skin microbiome in sensitive and resistant frogs and the data show a few remarkable distinctions between the three species studied. The presence of *Cladosporium, Elaphomyces, Monascus, Tritirachium, Ceratostomella* and *Claviceps* in *X. laevis* and their complete absence in *A. boreas* and *R. marina* is interesting because some of these fungi are reported to have novel anti-bacterial, anti-tumor (Hong, Seeram, Zhang, & Heber, 2008), toxin-producing (Zentmyer, 1942), or cuticle-degrading properties (Liang et al., 2010). However, none of them is known to be antagonistic to Bd (Table 1). Further, the presence of *Malassezia* in *A. boreas* and *R. marina* species and not in *X. laevis* is interesting because this fungus is known to cause skin infections in animals and humans and therefore may predispose these animals to Bd infections (Yim et al., 2010).

A number of hypotheses for Bd resistance in frogs have been postulated. Resistance to the fungus is likely the result of a combination of host and environmental factors and recent findings suggest that Bd resistance is not inborn. Voordouw and colleagues observed a stabilization in mortality rates due to Bd after five years of the pathogen’s introduction to British Columbia, but the population has yet to recover (Voordouw, Adama, Houston, Govindarajulu, & Robinson, 2010). This finding does not necessarily correlate to a corresponding decline in Bd incidence as suggested by Voordouw and colleagues (Voordouw et al., 2010). The distinction between a pathogenic dermatophyte and natural cutaneous microbiota is characterized less by the inherent properties of the organism and more so by the host’s ability to resist infection as suggested by Cogen *et al.* (Cogen et al., 2008).

From our results, we speculate that the skin microbiota plays a role in Bd resistance in *X. laevis* and *R. marina* frogs. In the case for *X. laevis*, this frog species had more diversity in fungal microbiome but not bacterial microbiome, and vice versa for *A. boreas*, we hypothesize that the lesser richness in fungal microbiota in *A. boreas* leave the host susceptible to Bd infection. Further, the poor fungal microbiome may be a consequent to the richness of their bacterial microbiome that can eliminate other fungi, leaving the host sensitive for Bd infection. Moreover, the high fungal diversity in *X. laevis* may protect against Bd infection due to the competition for nutrients or maybe due to the production of specific anti-Bd compounds. Finally, several factors could have influenced the skin microbiome for the tested frogs including their previous housing in either in a captive breeding facility, supply house, or housing facility.

## CONCLUSION

Numerous environmental and host-related factors can influence the susceptibility of frogs to Bd infection. One of which is the skin microbiota. We examined the skin microbiome of sensitive and resistant frogs to Bd and found unique differences in their bacterial and fungal microbiome. Investigating the role of the bacterial and fungal microbiota and their interactions are essential to elucidate frogs’ susceptibility to Bd infections and discover novel approaches to prevent or treat them by modulating the skin microbiota.

## METHODS

### Samples collection from frogs

Cotton tipped rods were soaked in a solution of 0.1 M sodium chloride and 10% Tween 20 then used to swab the adult animals on their ventral surfaces and thighs.

### Microbial diversity assessment with tag-encoded FLX amplicon pyrosequencing (bTEFAP)

Tag-encoded FLX amplicon pyrosequencing (bTEFAP) was performed as described previously at the Research and Testing Laboratory (RTL) Genomics (Lubbock, TX) (Dowd, Wolcott, Sun, et al., 2008). The new tag-encoded FLX-Titanium amplicon pyrosequencing (bTEFAP) approach is based upon similar principles but utilizes Titanium reagents and procedures and a one-step PCR, mixture of Hot Start and HotStar high fidelity taq polymerases, and amplicons originating from the 16S rRNA and ITS genes regions. The bTEFAP procedures were performed at the RTL Genomics (Lubbock, TX).

### DNA extraction

Total genomic DNA was extracted from the swab samples using a TissueLyser and QIAamp DNA mini kit per manufacturer directions (Qiagen, Velencia, CA) as previously described (Dowd, Callaway, Wolcott, et al., 2008). DNA samples were quantified using a NanoDrop spectrophotometer (Nyxor Biotech, Paris, France).

### Diversity data analysis

Following sequencing, all failed sequence reads, low quality sequence ends and tags were removed and sequences were depleted of any irrelevant sequences, to 16S rRNA or ITS, and chimeras using custom software and the Black Box Chimera Check software B2C2 as previously described (Dowd, Sun, Wolcott, Domingo, & Carroll, 2008; Gontcharova et al., 2010). Sequences less than 350 bp were removed. To determine the identity of taxa in the remaining sequences, sequences were first queried using a distributed BLASTn.NET algorithm against a database of high quality 16S rRNA bacterial sequences derived from the National Center for Biotechnology Information (NCBI) or ITS fungal sequences (Dowd, Zaragoza, Rodriguez, Oliver, & Payton, 2005). Database sequences were characterized as high quality based upon the criteria of RDP 9 (Cole et al., 2005). Using a .NET and C# analysis pipeline, the resulting BLASTn outputs were compiled, validated using taxonomic distance methods, and data reduction analysis performed as described previously (Acosta-Martínez, Dowd, Sun, & Allen, 2008). Rarefaction, Ace, and Chao1 to estimate mathematically predicted diversity and richness in the treatments using 350 bp trimmed, non-ribosomal sequenced depleted, chimera depleted, high quality reads was performed as previously described (Acosta-Martínez et al., 2008).

### Taxa identification

Based upon the above BLASTn derived sequence identity (percent of total length query sequence which aligns with a given database sequence), the taxa were classified at the appropriate taxonomic levels based upon the following criteria. Sequences with identity scores to known or well characterized sequences greater than 96.5% identity (i.e., < 3% dissimilarity) were resolved at the species level when possible, between 94.5% and 96.4% at the genus level, between 89.5% and 94.4% at the family and between 80% and 89.4% at the order level. After resolving based upon these parameters, the percentage of each taxa identification was individually analyzed for each sample providing relative abundance information within and among the samples based upon relative numbers of reads within a given sample. When multiple identifications were found, such identifications were resolved arbitrarily to the top hit. Evaluations presented at a given taxonomic level, except species level, represent all sequences resolved to their primary genera identification or their closest relative, where indicated.

### Statistical analyses

Double dendrograms were performed using comparative functions and multivariate hierarchical clustering methods of NCSS 2007 (NCSS, Kaysville, Utah) based upon the most abundant genera, Weighted-pair group clustering method, and Manhattan distance method with no scaling. Manhattan distance for the relative percentage data is calculated between rows *j* and *k* using:

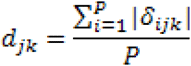

where *δ_ijk_* = *z_ij_* – *z_jk_*

It should be noted that the dendrogram linkages of the classes are not phylogenetic but based upon abundance of genera among the samples ordered in rows. Clustering of the systems were similarly based upon abundance of the top 40 most abundant genera among individual samples.

## DATA AVAILABILITY

All data reported in this study are included in this manuscript.

